# Computational Modeling of Motor-Driven Extension of Microtubule Bundles in Axons

**DOI:** 10.1101/2025.09.25.678704

**Authors:** Donghyun Yim, Laurel Patterson, Daniel Suter, Kyle Miller, Taeyoon Kim

## Abstract

In addition to actin assembly at the growth cone, neuronal axons elongate via interactions between microtubules and microtubule-associated proteins including dynein motor proteins and cross-linking proteins. Dynein translocates microtubules toward the growth cone to exert extensile forces for axonal outgrowth. During this process, microtubules are most likely to experience compressive loads, which can result in bending deformation on microtubules called buckling. Such buckled microtubules are impaired in their ability to bear compressive forces and may not contribute significantly to axonal outgrowth. If microtubules are interconnected by cross-linking proteins, they are less likely to be buckled and thus able to resist larger compressive loads. Despite the importance of the buckling and connectivity of microtubules, their effects on the axonal outgrowth have not been investigated to date. In this study, using an agent-based computational model, we created a microtubule bundle to simulate the microtubule system in axons. This bundle increased its length by interactions between dynein-like motors and microtubules against a mechanical load. We found that intermediate cross-linking density resulted in maximal elongation of the bundle, because microtubules were easily buckled at low cross-linking density, whereas the displacement of microtubules was inhibited at high cross-linking density. When individual microtubules were much stiffer, the bundle showed higher elongation even with lower cross-linking density, since stiff microtubules were less likely to be buckled by compressive loads. Our study provides new insights into the mechanisms driving axonal outgrowth.

## INTRODUCTION

Neurite outgrowth is a process in which one or more projections extend from the cell body of neurons. Only one of the projections elongates fast and develops into axons, whereas the rest becomes dendrites after much slower extension ^1^. Outgrowth of the axons guided by physical and chemical cues creates a complex neural network by forming synaptic connections between neurons. Axonal outgrowth is known to be driven mainly by molecular interactions between microtubules (MTs), microtubule-associated proteins (MAPs) ^2-4^, and actin ^5^. MTs are biopolymers made of α- and β-tubulin dimers with a hollow cylinder structure of 25 nm in diameter and with polarity defined by plus and minus ends ^6^. In case of cultured hippocampal neurons, there are ∼1,400 MTs with 5 µm in length on average in a stage 3 axon, and MTs can elongate up to tens of µm ^7^. In higher organisms, the plus ends of MTs in axons are directed toward axon terminals, unlike the random orientation of MTs within dendrites ^8^. These MTs are initially polymerized at centrosome and then carried into and along axons by MT-motor proteins called kinesin and dynein ^4,9-11^. These motor proteins are well-known for mediating axonal transport of cargo along MTs. Kinesin walks along MTs toward their plus end, thus delivering cargos, such as vesicles, proteins, RNA, and organelles, toward the axon terminal during anterograde transport ^12^. Conversely, dynein walks toward the minus end of MTs, thus mediating retrograde transport in axons ^13^. In addition to these motor proteins, MT cross-linkers also play an important role in physically interconnecting MTs ^14^. In axons, Tau is known as a primary cross-linker with a highly transient binding behavior characterized by frequent binding and unbinding ^15,16^.

It has been hypothesized that dynein bound to the plus end of one MT displaces the adjacent MT by walking toward the minus end of this MT, which results in MTs being translocated toward the axon terminal to drive axonal outgrowth ^17-20^, demonstrated by several previous studies on the crucial role of stable MTs and dyneins ^21-25^. In addition, axonal outgrowth is also driven by the growth cone which steers and guides the neurite to its destination by actomyosin-based contraction ^26,27^. It was shown that the disruption of actin filaments at growth cone inhibited the axonal elongation ^28^. However, these forces originating from MTs, dyneins, and the growth cone are partially counterbalanced by contractile forces generated by actomyosin cortex enveloping MTs in the axons. The actomyosin cortex in the axons consists of an array of separate actin rings interconnected by spectrin dimers ^29-32^ and myosin IIA along the axonal shaft suppress neurite elongation by generating contractile forces to the axon ^27,33,34^.

To understand the mechanism of axonal outgrowth, various computational models have been developed. These models comprised discrete MTs, cross-linkers, and dynein motors within a confining computational domain ^35,36^, revealing the contributions of dynein motors and cross-linkers to the extension of the MT bundle. However, all of these former models did not account for the deformation of MTs induced by mechanical forces; MTs were simplified into infinitely rigid straight segments moving only in the axial direction of neurites. Thus, the consideration of volume exclusion between neighboring MTs was not necessary in these models. Although MTs are much more rigid than other cytoskeletal polymers, they can be bent and buckled if they feel compressive forces greater than a critical buckling force ^37^. Indeed, it has been shown that MTs in cells can exhibit high curvatures ^38-42^. During axonal elongation, MTs must experience large compressive loads when they exert extensile forces against the actin cytoskeleton and plasma membrane at the axon terminal. Therefore, it is of great importance to properly represent the deformability of MTs.

To overcome these limitations and understand how deformable MTs mediate axonal outgrowth, we developed an agent-based model for a bundle consisting of MTs, cross-linkers, and dynein-like motors within an axon-like long cylindrical space. This MT bundle can elongate via interactions between motors and MTs described earlier against a resisting force acting on one end of the cylindrical space. Using this model, we probed how the connectivity and rigidity of MTs affect the extensile force generated by the MT bundle. Note that although our system is inspired by axonal outgrowth, this bundle model does not completely replicate the axon structure.

## MATERIALS AND METHODS

In our model, all elements constituting a bundle––MTs, cross-linkers, and motors––are confined within a cylindrical domain whose radius and length are 2 µm and 20 µm, respectively (Fig. 1A). One circular boundary of the domain is fixed, whereas the opposite boundary can move to increase the length of the domain if the bundle exerts pushing forces greater than a resisting force, *F*_*R*_. The moving boundary represents the axon terminal or growth cone. MTs, cross-linkers, and motors are simplified via cylindrical elements; MTs are coarse-grained into serially-connected cylindrical elements. Cross-linkers, simplified into two cylindrical elements connected at their center, transiently interconnect MTs. Motors, which also consist of two cylindrical elements, displace MTs toward the moving boundary.

**Fig. 1.**
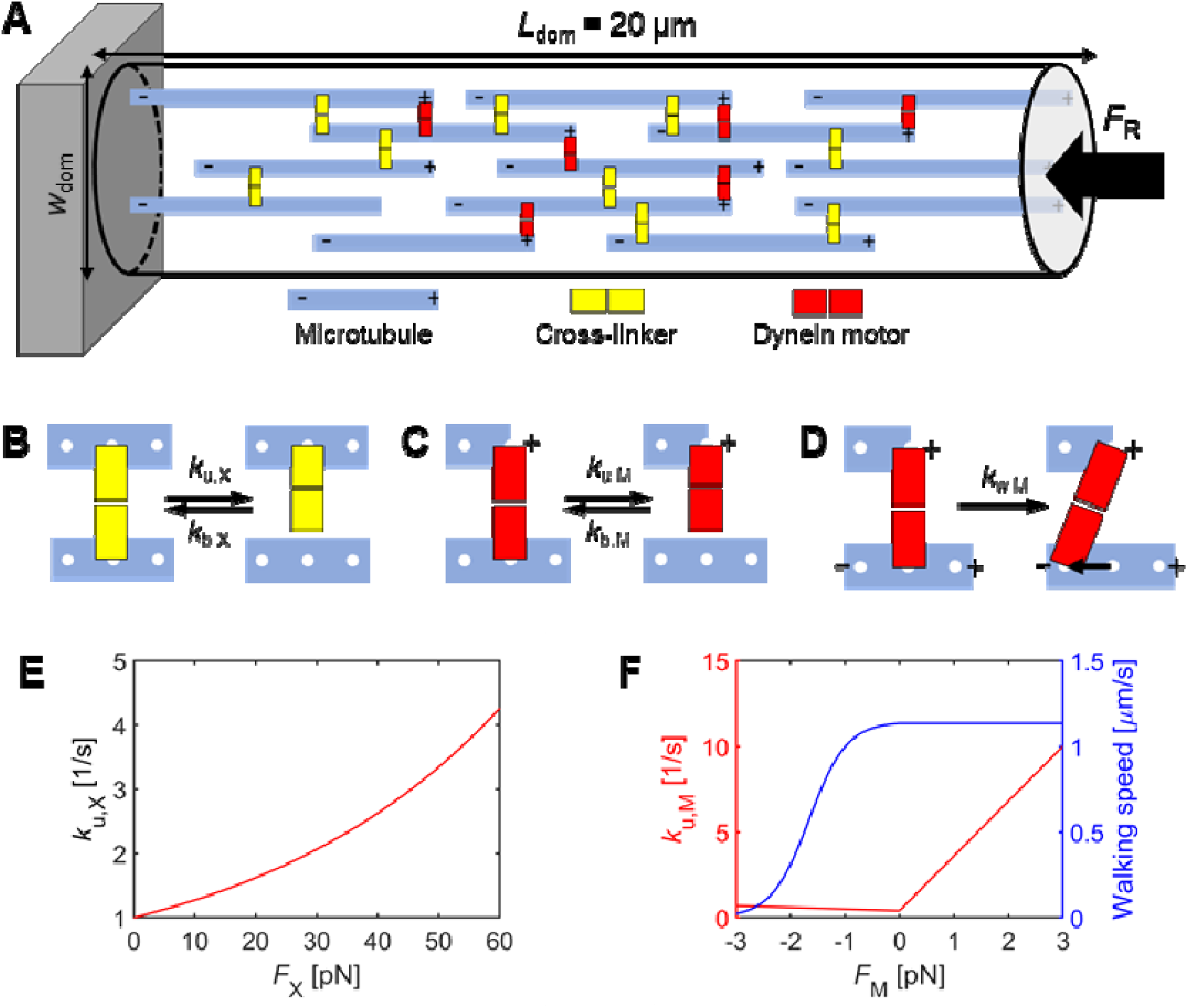
Agent-based computational model. (A) A bundle consisting of microtubules (MTs, blue), cross-linkers (yellow), and dynein-like motors (red). MTs are distributed in random positions with different lengths within a cylindrical space whose diameter (thickness) and initial length are *w*_dom_ and *L*_dom_, respectively. All of their plus ends are initially directed toward the extending end which exerts a resisting force to the bundle (). The minus ends of some of the MTs are clamped on the opposite stationary boundary. *w*_dom_ is 0.6 μm in most of the simulations, and *L*_dom_ increases over time as MTs push the extending end by overcoming the resisting force. (B-D) Dynamic behaviors of cross-linkers and motors. (B) Cross-linkers bind to MTs with a constant rate, *k*_b,X_, and unbind from MTs at a force-dependent rate, *k*_u,X_. (C) The cargo end of motors is permanently bound to the plus end of MTs. The walking end of motors binds to any part of MTs at a constant rate,, and unbinds from MTs at. (D) The walking end walks toward the minus end at a rate,, by shifting its binding site by ∼8 nm. (E) The dependence of on a tensile force applied on a cross-linker (*F*_X_). (F) (red) and walking speed (blue) are determined by a force that a motor feels (). Note that the walking speed is equal to ×8 nm. *F*_M_ is considered positive if a motor is pulled toward the plus end of MTs.

### Brownian dynamics

At each time step, the velocity of the nodes of all cylindrical elements is calculated by the

Langevin equation:

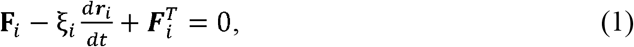

where **F**_*i*_, ***r***_*i*_, and *ξ*_*i*_ represent the deterministic force, position, and drag coefficient of *i*^*th*^ element, respectively, and *t* is time. 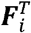 is a stochastic force satisfying the fluctuation-dissipation theorem ^43^:

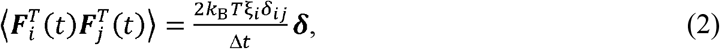

where *δ*_*ij*_ is the Kronecker delta, ***δ*** is a second-order tensor, *k*_B_*T* is thermal energy, and Δ*t* = 1.8e-5 s is time step. *ξ*_*i*_ is calculated using the following approximated form for a cylindrical object ^44^:

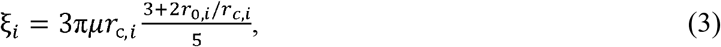

where *μ* is the viscosity of a surrounding medium, and *r*_0,*i*_ and *r*_*c,i*_, are the equilibrium length and diameter of an element, respectively. ***r***_*i*_ is updated via the velocity, d ***r***_*i*_ /d*t*,, calculated from Eq. 1 and via the forward Euler scheme:

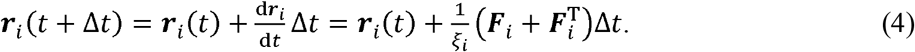

The deterministic force,***F***_*i*_, includes extensional, bending, and repulsive forces. The extensional and bending forces originate from the following harmonic potentials:

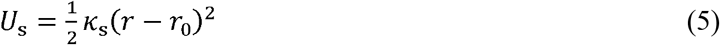

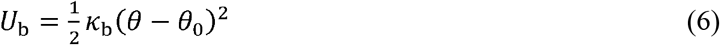

where *K*_s_ and K_b_ indicate the extensional and bending stiffnesses, *r* is the instantaneous length, and *θ* and *θ*_0_ are instantaneous and equilibrium angles. An equilibrium angle formed by two adjacent MT elements (*θ*_0,MT_ = 0 rad) and the equilibrium length of each MT element (*r*_0,MT_ = 280 nm) are maintained by bending (*κ* _b,MT_) and extensional (*κ* _s,MT_) stiffnesses of MTs, respectively. The reference value of 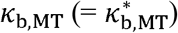 corresponds to the persistence length of 1 mm ^45^. An equilibrium angle formed by two elements of a cross-linker (*θ*_0,X1_ = 0 rad) and that formed by one element of the cross-linker and a MT element where the cross-linker is bound (*θ*_0,X2_ = π/2 rad) are maintained by two bending stiffnesses (*κ*_b,X1_ and *κ*_b,X2_). The equilibrium length of each cross-linker element (*θ*_0,X1_ = 20 nm) is maintained by the extensional stiffness of cross-linkers *(κ* _s,X_). The equilibrium length of each motor element (*r*_0,M_ = 10 nm) is maintained by the extensional stiffness of motors (*r*_s,M_). There is no bending stiffness involved with motors.

Volume-exclusion effects between overlapping MT elements are considered via repulsive forces originating from the following potential:

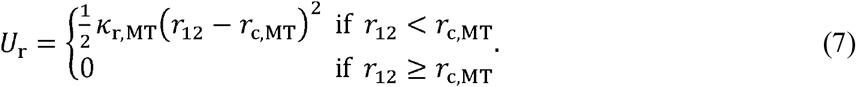

where *κ*_r,MT_ is the strength of the volume-exclusion effects, *r*_12_ is a minimal distance between MT elements, and *r*_c,MT_ is a threshold distance which is equal to the diameter of MTs.

Forces acting on MT elements due to bound cross-linkers and motors or due to repulsive forces are distributed onto the two nodes of the MT elements as described in our previous work in detail ^46^; as the node is closer to the point of force application, it feels a larger portion of the force.

### Dynamic behaviors of MTs, cross-linkers, and motors

MTs are assembled in a stochastic manner via nucleation and polymerization. Simulations begin with the monomer pool of MT elements. First, for the nucleation event, one MT cylindrical element is drawn from the pool and appears in space as an explicit element. Then, this element is polymerized at the plus end via the addition of additional cylindrical elements. For simplicity, it is assumed that MTs do not undergo any type of disassembly as in previous models ^35,36^.

Considering the size of tubulin dimers (∼8 nm) and 13 columns in the MT lattice structure, there are 35-equispaced binding sites on each MT element, and up to 13 cross-linkers or motors can bind to one binding site simultaneously. Each cross-linker, consisting of two interconnected cylindrical elements, has two free ends. The free ends of cross-linkers bind to binding sites on MTs at a constant rate (*k*_b,x_) and unbind from MTs at a force-dependent rate (*k*_u,x_) determined by the Bell’s equation (Fig. 1B) ^47^:

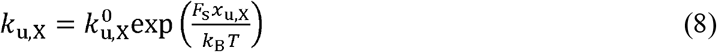

where 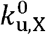 is a zero-force unbinding rate constant (Fig. 1E). *F*_*s*_ and *x*_u,x_ are the magnitude of a spring force acting on the cross-linker element and force sensitivity, respectively.

Two cylindrical elements constituting each motor represent “cargo” and “walking” domains. The free end of the cargo element permanently binds to the plus end of MTs. It is assumed that only one motor is allowed to bind to each plus end to avoid the saturation of the plus ends by too many motors. The free end of the walking element can bind to the other MT at a constant rate (*k*_b,M_), walk toward the minus end at a force-dependent rate (*k*_w,M_), and unbind from the MT at a force-dependent rate (*k*_u,M_) (Figs. 1C, D). The walking motion is implemented by displacing the binding site of the walking element by ∼8 nm at each walking event. The force dependence of *k*_u,M_ and *k*_w,M_ are adopted from a previous study (Fig. 1F) ^48^.

### Quantification of MT curvature

To identify curved MTs, the curvature of MTs is calculated as follows. For MT consisting of *N* nodes, the curvature is computed on all nodes except two nodes located at plus and minus ends (1< *i* < N). On each node *i*, two vectors are calculated, ***a***_*i*_ = ***r*** _*i* + 1_ − ***r***_*i*_ and *b*_*i*_ = ***r***_*i*_ − ***r*** _*i* −1_, where ***r*** _*i* −1_, ***r***_*i*_, and ***r*** _*i* + 1_ are the positions of three adjacent nodes. Then, the curvature is calculated as:

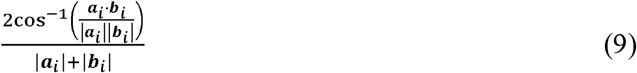

If the curvature is greater than 0.2 rad/µm, it is assumed that this part of MT on the node *i* experiences significant bending deformation ^49^.

### Energy calculation

Due to interactions between elements described earlier, elastic energy is developed on each element. Extensional (i.e., spring) and bending energies stored in MTs, cross-linkers, and motors are quantified. Note that tensile and compressive energies, both of which are extensional in nature, are separately measured depending on whether an instantaneous length is greater or

smaller than an equilibrium length. In case of cross-linkers, average extensional energy for their two elements is measured, and two types of bending energies involved with *κ*_b,X1_ and *κ*_b,X2_ are summed. In the case of motors, only average extensional energy for their two elements is considered.

### Simulation procedures

All simulations start with the bundle formation phase for 1 s. During this phase, monomeric MT elements undergo nucleation and polymerization events, resulting in the formation of MTs in random locations with their plus ends all directed in +z direction (i.e., toward the moving circular boundary of the cylindrical domain). After the depletion of all MT elements, the average length of MTs becomes ∼7 μm under the reference condition. In addition, cross-linkers and motors bind to MTs during this phase. Note that motors do not walk during the bundle formation phase. After the bundle is formed, motors start walking toward the minus end of MTs, and various quantities are measured for 1 hr.

### Statistical analysis

Two-tailed unpaired t-test is conducted using MATLAB 2021b. Sample size is provided in figure captions.

## RESULTS

### Extension of microtubule bundles without cross-linkers can be modulated by several factors

First, to clearly understand the role of dynein activity in bundle elongation, we performed a simulation under a reference condition without any cross-linker for 1 hr (Fig. 2) and then compared results from the simulations with those reported in previous computational studies that also excluded cross-linkers ^36,50^. It was observed that bundle extension level (*ε*) became maximal at ∼100 s and then decreased afterwards, and highly curved MTs were observed near the extending end (Figs. 2A, B). For the first ∼20 s, compressive energy measured on all MTs decreased, whereas their bending energy increased, and then they hardly changed after ∼20 s (Fig. 2C). The fraction of curved MTs also increased during for first ∼20 s of the bundle extension (Fig. 2D), consistent with the increase in the bending energy. Most of these curved MTs were found near the extending end (Fig. S1). The increased curvature of MTs arises from interactions between MTs and motors and also between MTs and surrounding environments (i.e., the boundaries of the computational domain). Considering the decreased compressive energy, it is likely that compression-induced buckling was the main reason for the increased curvature of MTs. The increase in the number of buckled MTs could explain the decrease in *ε* at later times. However, it is possible that motor walking activity might have weakened at later times. To check this possibility, we measured average motor walking speed and average tension acting on motors (Figs. 2E, F). It was found that motors maintained walking speed and tension even after reaching their peak levels, negating the contribution of motor impairment to reduced *ε* at later times.

**Fig. 2.**
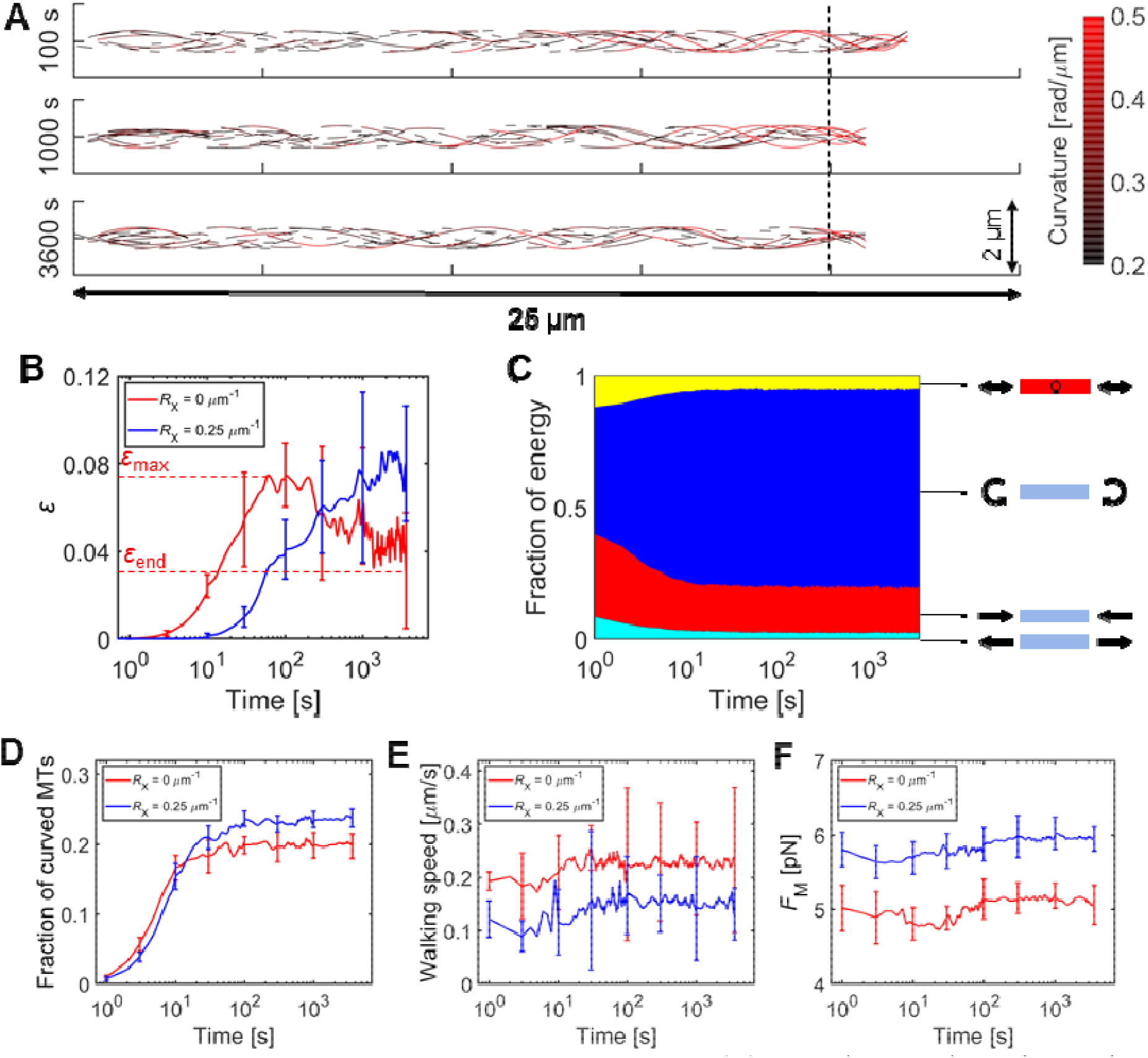
Bundle extension with or without cross-linkers. (A) Snapshots at three time points showing a fraction of microtubules (MTs) with curvature greater than 0.2 rad/µm. Only MTs were visualized for clarity. 10 simulations were run for 1 hr under this condition. MTs near the extending end on the right tend to be curved more. (B) Time evolution of normal strain (*ε*) calculated by dividing an increase in the domain length (*L*_dom_) by the initial length, 20 µm. When *R*_X_ = 0 µm^-1^, after reaching peak level (*ε*_max_), *ε* gradually decreased and reached final level (*ε*_end_) at 1 hr. When *R*_X_ = 0.25 µm^-1^, *ε* did not change significantly over time after ∼1,000 s. (C) The fraction of four types of energies over time: tensile (cyan), compressive (red), and bending (blue) energies of MTs and the extensional energy of motors (yellow). (D) The fraction of curved MTs over time. (E) The walking speed of motors and (F) forces acting on motors (*F*_M_) over time. Motor activities did not show a large change after ∼100 s. Note that cross-linkers were excluded in (A) and (C). Their counterparts with cross-linkers are Figs. S2A, B.

Then, we repeated simulations with a variation in each of the parameters which are known to affect bundle extension — tubulin concentration (*C*_MT_), average MT length (*L*_MT_), motor density (*R*_M_), and domain thickness (*w*_dom_) — and measured strain level at the final time point (*ε*_end_), with cross-linkers still excluded (Fig. 3). Under the reference condition used earlier, the following parameter values were employed: *C*_MT_ = 23 µM, *L*_MT_ = 7 µm, *R*_M_ = 5 µm^-1^, and *w*_dom_ = 0.6 µm. In previous studies, it was reported that a larger number of MTs or dynein motors enhanced the bundle extension, whereas an increase in the average length of MTs suppressed the bundle extension ^50-52^. In our simulations, the bundle could not elongate (i.e., *ε*_end_ ∼ 0) when *C*_MT_ was lower than 20 µM (Fig. 3A), which could be partially attributed to lack of percolation between two circular boundaries of the computational domain. Above 20 µM, *ε*_end_ increased almost linearly as *C*_MT_ increased. Since we fixed *R*_M_ (= *N*_M_ /∑*L*_MT_ where *N*_M_ is the number of motors), higher *C*_MT_ also increased the total number of motors. Then, the bundle with more MTs and more motors can naturally exert a stronger extensile force to the moving boundary of the domain, resulting in higher bundle extension. When MTs were longer on average (i.e., higher *L*_MT_), *ε*_end_ tended to be greater in general, but at very high *L*_MT_, *ε*_end_ slightly reduced (Fig. 3B). This is because longer MTs can be pushed toward the domain boundary by more dynein motors, but there would be fewer plus ends where motors can be anchored via the cargo end. Higher *R*_M_ showed a positive effect on *ε*_end_, but as *R*_M_ increased more, the rate of an increase in *ε*_end_ became lower (Fig. 3C). This is attributed to the limited number of plus ends; if most of the plus ends are already saturated with motors, a further increase in the number of motors would not enhance bundle extension significantly. If more than one motor is allowed to bind to each plus end in this model, it is expected that the bundle extension would be sensitive to a change in the number of motors even at high level of *R*_M_. Lastly, when *w*_dom_ was varied, *ε*_end_ was inversely proportional to *w*_dom_ because a narrower domain can prevent MTs from undergoing too much curvature by supporting MTs from their side (Fig. 3D). When *w*_dom_ became greater than ∼0.7 µm, *ε*_end_ was negligibly small due to lack of side support for MTs. Note that the reference domain thickness, 0.6 µm, is close to the thickness of MT bundles found in neuronal axons ^53^. Overall, our results obtained without cross-linkers showed good qualitative agreements with previously reported data in the literature in terms of the effects of the key parameters on bundle extension.

**Fig. 3.**
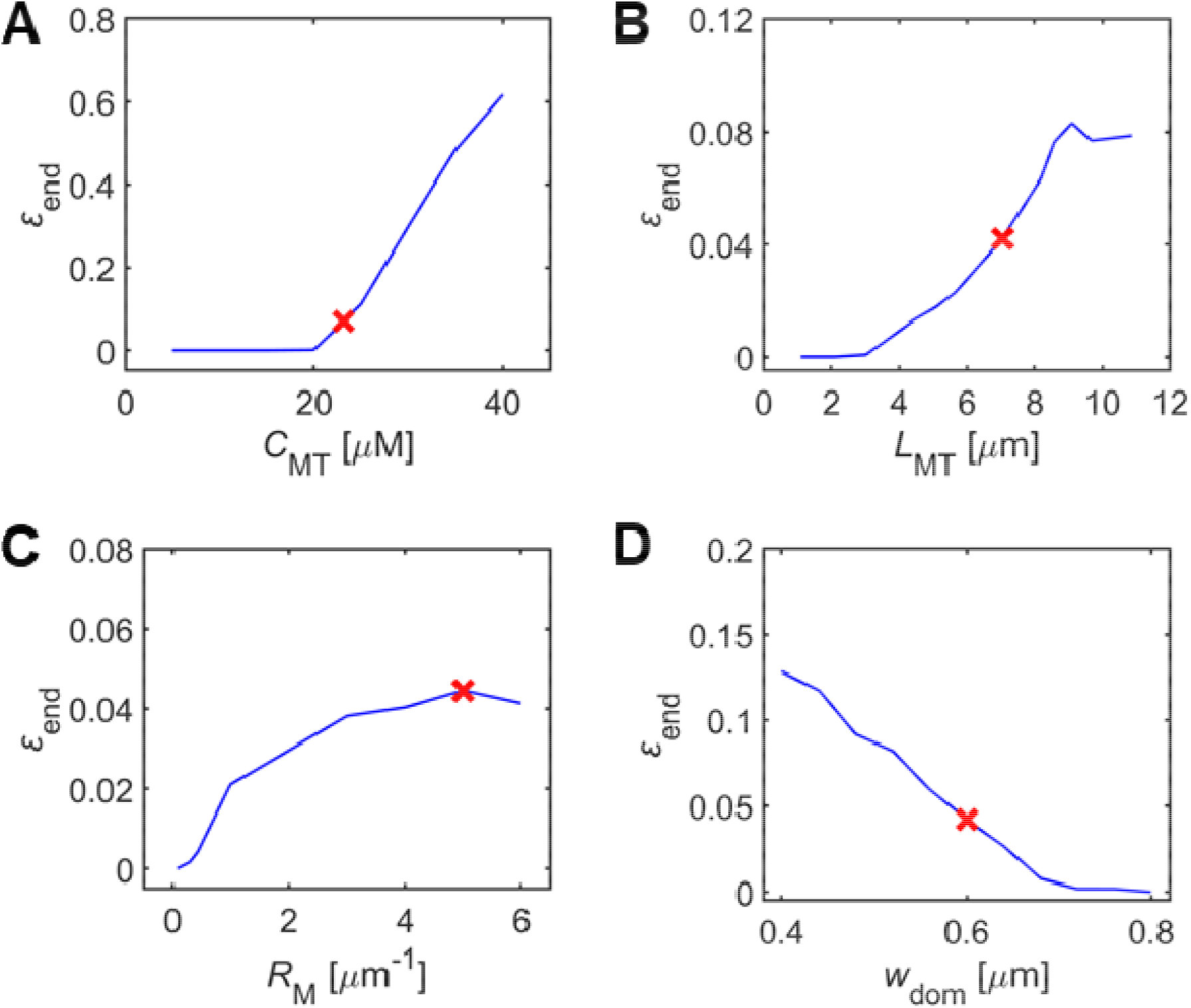
Effects of four parameters on the final level of bundle extension (*ε*_end_) without cross-linkers. (A) Concentration of microtubules (MTs) (*C*_MT_) (*n* = 10), (B) average length of MT (*L*_MT_) (*n* = 20), (C) motor density (*R*_M_) (*n* = 50), and domain thickness (*w*_dom_) (*n* = 10). A red cross in each panel indicates the reference case shown in Fig. 2 with, *C*_MT_ = 23 µM, *L*_MT_ = 7 µm, *R*_M_ = 5 µm^-1^, and *w*_dom_ = 0.6 µm.

### Bundle extension showed biphasic dependence on cross-linking density

Cross-linkers are known to play a crucial role in stabilizing MTs in neuronal axons by interconnecting MTs ^54,55^. It was hypothesized that more anchoring points help MTs withstand larger compressive loads by increasing a minimum force required for buckling ^37^, which would enhance maximal bundle extension level (*ε*_max_) and *ε*_end_. However, cross-linkers also increase effective friction between MTs, which makes the displacement of MT relative to other MT harder. To understand the effects of cross-linkers on bundle extension, we investigated the case in presence of cross-linkers at *R*_X_ = 0.25 µm^-1^ (0.25 cross-linkers per μm along MTs on average). Notably, the bundle reached *ε*_max_ much slower at ∼1,000 s than the case without cross-linkers (Figs. 2B and S2A). However, unlike the bundle without cross-linkers, *ε* remained steady until ∼ 3,600 s. The tendency of changes in bending and compressive energies was similar to that observed without cross-linkers; compressive energy decreased, whereas bending energy increased for first ∼20 s (Fig. S2B). The fraction of curved MTs reached peak at ∼20 s and remained at similar level (Fig. 2D). Interestingly, the peak level was slightly higher than that without cross-linkers. With cross-linkers, motor walking speed was lower (Fig. 2E), and tension acting on motors was significantly higher than without cross-linkers (Fig. 2F). Without cross-linkers, motors can displace MTs without a significant resistance. By contrast, if cross-linkers exist on MTs, motors should exert stronger forces to achieve the same displacement of MTs, resulting in slower motor walking. The higher fraction of curved MTs would originate from buckling induced by a competition between motors and cross-linkers, which can occur on any point of MTs (Fig. S3). Indeed, it was observed that more MTs were curved in regions away from the extending end (Fig. S2C).

Then, we repeated simulations with a variation in cross-linking density (*R*_X_) between 0.01 µm^-1^ to 10 µm^-1^ and then measured *ε*_max_ between 0 s and 1 hr and *ε*_end_ at 1 hr. As *R*_X_ was reduced from 0.25 µm^-1^ used for the reference case, a bundle reached slightly higher *ε*_max_ more quickly, but *ε*_end_ significantly decreased, which means that bundle extension was less sustainable with lower *R*_X_ (Figs. 4A and S4A-B and Movies S1-S3), as in the case without cross-linkers (Fig. 2B). When *R*_X_ was increased from 0.25 µm^-1^, both *ε*_max_ and *ε*_end_ substantially decreased. *ε*_end_ was similar to *ε*_max_ at higher *R*_X_, indicating that bundle extension was sustained after reaching its peak level or was still increasing at 1 hr. Indeed, time for *ε*_max_ was greater than 1,000 s when *R*_X_ ≥ 0.1 µm^-1^. (Fig. S4C). Overall, *ε*_end_ showed biphasic dependence on *R*_X_ with its maximum at the reference value of *R*_X_ = 0.25 µm^-1^, implying the existence of two distinct factors that obstruct bundle extension at extremely low and high *R*_X_.

**Fig. 4.**
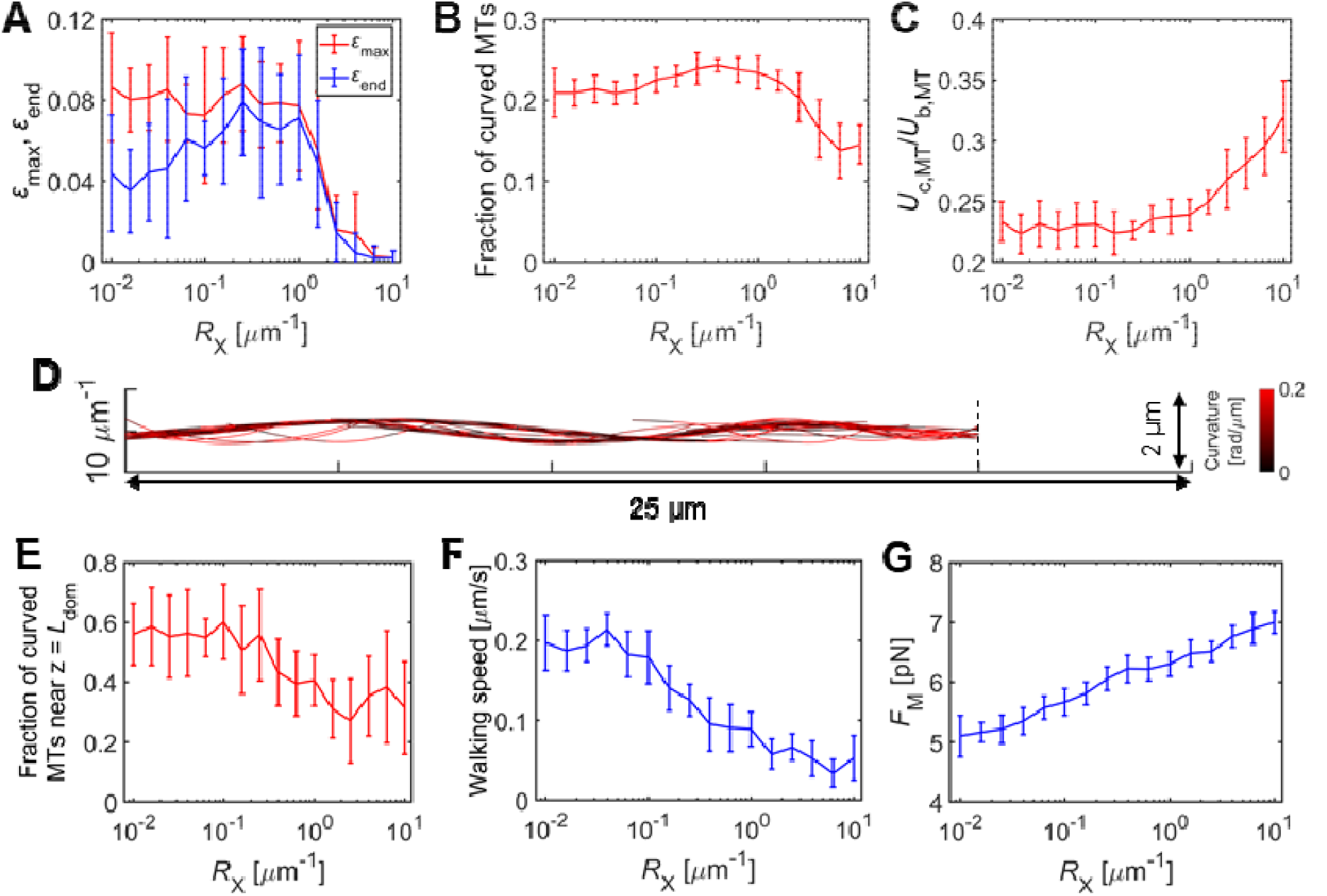
Bundle extension depending on cross-linking density (*R*_X_). 10 simulations were run with each value of *R*_X_. (A) Maximal normal strain (*ε*_max_) and final normal strain (*ε*_end_) identified between 0 s and 3,600 s with different *R*_X_. (B) The fraction of curved microtubules (MTs) and (C) the ratio of MT compressive energy to MT bending energy as a function of *R*_X_. (D) Snapshot of MTs in the bundle with their curvature represented via color scaling with a large number of cross-linkers (*R*_X_ = 10 µm^-1^). (E) The fraction of MTs located near the extending end with different *R*_X_. (F) The average walking speed of motors and (G) forces acting on motors () depending on *R*_X._

We found that one of the two factors, which affects bundle extension as a function of *R*_X_, is related to MT bending. As *R*_X_ increased from 0.25 µm^-1^, the fraction of curved MTs was substantially reduced, and the ratio of compression energy to bending energy in MTs elevated (Figs. 4B, C). Euler’s buckling load predicts that a critical compressive load for buckling is proportional to the bending stiffness of MTs (*κ*_b,MT_) and inversely proportional to the square of the length of an unsupported part (*L*_uns_):

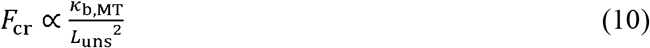

If MTs are constrained by cross-linkers, *L*_uns_ corresponds to spacing between adjacent cross-linking points. As *R*_X_ increases, *L*_uns_ decreases on average, increasing *F*_cr_ and thus reducing the frequency of buckling. If *R*_X_ becomes very large, most of the MTs behave as a single entity that has much higher effective bending stiffness (Fig. 4D). As *R*_X_ decreased from 0.25 µm^-1^, the fraction of curved MTs was slightly reduced to level observed in the case without any cross-linker (Fig. 4B). As explained earlier, this is related to buckling induced by the competition between motors and cross-linkers (Fig. S3). With fewer cross-linkers, motor-induced buckling, which occurs independently of interactions with the extending end, was less likely to take place, so the fraction of curved MTs was reduced. However, the fraction of curved MTs near the extending end was inversely proportional to *R*_X_ in general (Fig. 4E), and the orientations of the plus ends of these MTs deviate more from the axial direction (Fig. S4D). These imply that MTs can push the extending boundary better with more cross-linkers by resisting buckling. The other factor affecting bundle extension is the mobility of MTs. Transient cross-linkers result in effective friction between MTs ^56^. As the number of cross-linkers increases, MTs are harder to be displaced by motors (relative to other MTs) due to large friction, which leads to slower bundle extension. With higher *R*_X_, motors experienced larger forces and walked more slowly due to such a resistance from cross-linkers (Figs. 4F, G). In sum, the bundle extension is optimal at intermediate *R*_X_ because by two governing factors of bundle extension––how strongly MTs can resist compressive loads and how fast MTs can be displaced––show opposite dependence on *R*_X_.

### Stiffness of MTs has a large effect on bundle extension

As shown in Eq. 10, the critical load for buckling (*F*_cr_) is proportional to *κ*_b,MT_. Thus, if individual MTs becomes stiffer, MTs would be able to support higher compressive loads. We probed the effect of *κ*_b,MT_ on bundle extension in the presence of cross-linkers (*R*_X_ = 0.25 μm^-1^) by increasing the persistence length of MTs (*l*_p_), which is proportional to *κ*_b,MT_:

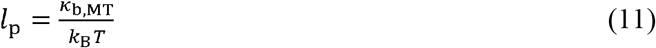

Note that *l*_p_ in all previous results was 1 mm. As expected, higher *l*_p_ increased *ε*_end_ (Fig. 5A and Movies. S4, 5). The lower curvature of MTs and the higher ratio of compressive energy to bending energy with greater *l*_p_ imply that MTs were less prone to buckling when they attempted to push the extending end (Figs. 5B, C and S5A). Indeed, MTs near the extending end were less curved and deviated less from the axial direction (Figs. 5D, E and S5B). Unlike the case with *l*_p_ = 1 mm (Fig. S2C), MTs showed relatively uniform curvature regardless of their axial (i.e., *z*) position (Fig. S5C). The change in *l*_p_ did not significantly vary the average walking speed of motors (Fig. S5D), meaning that enhanced *ε*_end_ was not associated with a change in motor activities.

**Fig. 5.**
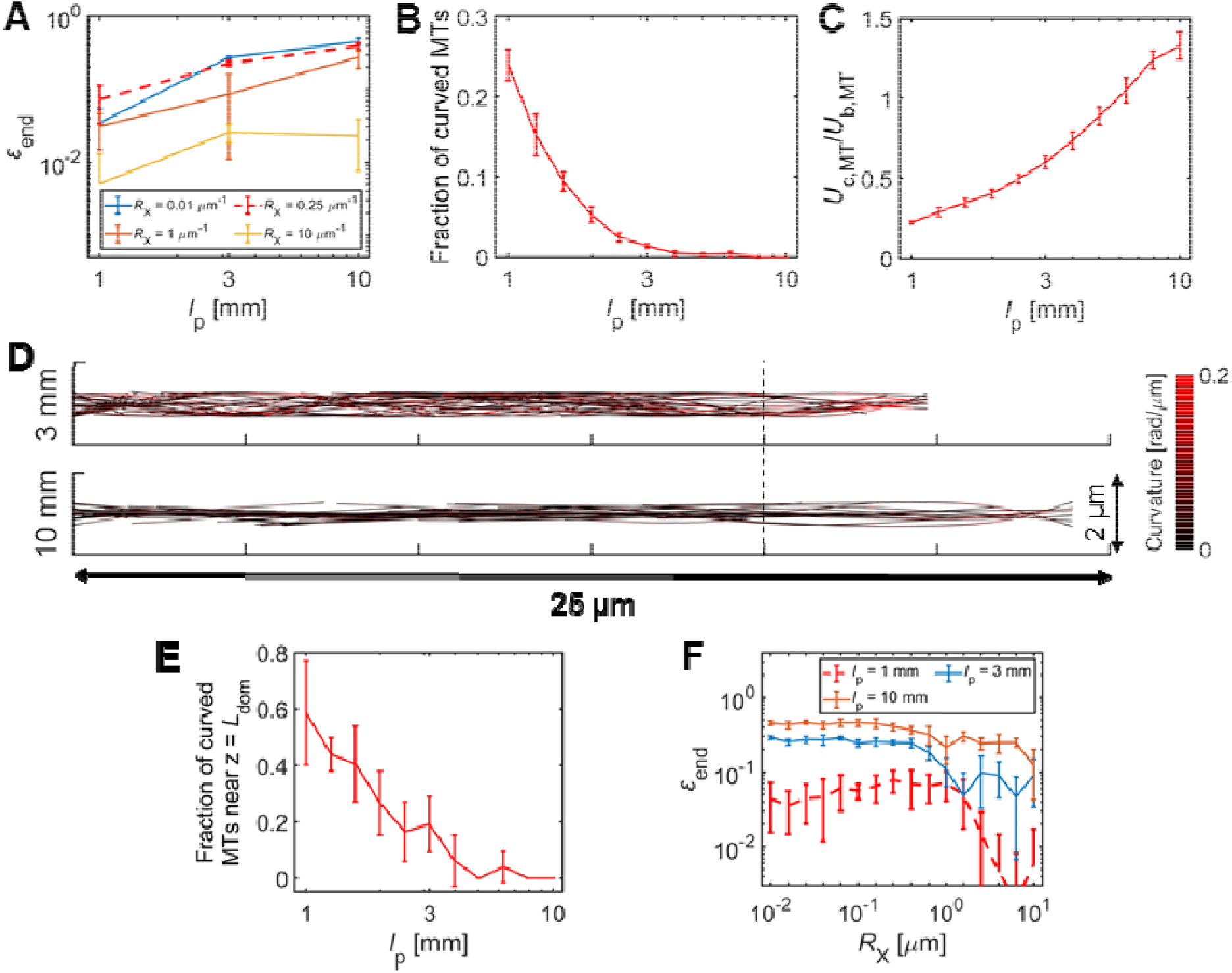
Bundle extension with different persistence length of microtubules (*l*_p_). 10 simulations were run for each value of *l*_p_. (A) Final normal strain measured at 1 hr (*ε*_end_) as a function of *l*_p_ with four different *R*_X_. (B) The fraction of curved MTs and (C) the compressive-to-bending energy ratio of MTs. (D) Snapshots of two cases with *l*_p_ = 3 mm (top) and 10 mm (bottom) with MT curvature visualized via color scaling. *R*_X_ was 0.25 µm^-1^ in these two cases. (E) The fraction of curved MTs located near the extending end with various *l*_p_. (F) The dependence of *ε*_end_ on *R*_X_ with 3 different *l*_p_. The biphasic dependence disappeared with higher *l*_p_.

Considering that cross-linkers on MTs affect *F*_cr_ (Eq. 10), it is expected that the effect of *l*_p_ would depend on *R*_X_. Thus, we repeated simulations with various values of *l*_p_ and *R*_X_ (Figs. 5A, F). It was found that when cross-linking density was very high (*R*_X_ = 10 μm^-1^), *ε*_end_ did not increase significantly with higher *l*_p_ because the restriction of MT movement induced by a large number of cross-linkers played a dominant role in determining *ε*_end_ (Fig. 5A). Interestingly, with high *l*_p_ (= 3 mm or 10 mm), the biphasic dependence of *ε*_end_ on *R*_X_ disappeared (Fig. 5F); *ε*_end_ was largely independent of a variation in *R*_X_ at *R*_X_ < 0.25 μm^-1^ because MTs were already stiff enough to resist compressive loads unlike MTs with *l*_p_ = 1 mm. However, even with such stiff MTs, *ε*_end_ decreased at *R*_X_ ≥ 0.25 μm^-1^ because the restriction of MT movement, which plays a dominant role for *ε*_end_ at high *R*_X_, is not related to how stiff MTs are. The average walking speed of motors in the case with high *l*_p_ did not show a noticeable difference (Fig. S5E), implying that the change in the dependence of *ε*_end_ on *R*_X_ did not originate from different motor activities.

## DISCUSSION

During axonal outgrowth, molecular interactions between MTs, cross-linkers, and dynein motors generate extensile forces required for elongating axons ^3^. Previous experiments revealed that dynein motors translocate MTs toward axon terminals to extend axons ^19,20,57^, and that Tau cross-linkers is required to ensure the axonal elongation ^2,58^. The extensile forces are partially counterbalanced by contractile forces generated by actomyosin envelopes ^59^. Several computational studies have demonstrated how these mechanical forces are generated and compete with mechanical resistances for outgrowth ^35,36^ or how a cross-linked MT bundle resists the compressive force ^60^. However, none of these models considered both, so how deformability of MTs is related to neuronal outgrowth remains elusive, although they can undergo buckling by compressive loads from surrounding structures. Motivated by the axonal outgrowth process and one of the previous models ^36^, we developed a computational model to investigate how bundle extension is regulated by deformable MTs and other key factors. In our model, MTs interconnected via cross-linkers are translocated by dynein-like motors to push one boundary of the computational domain where a resisting force is applied.

First, we performed simulations without any cross-linker to test the effects of four parameters—the concentration and length of MTs, motor density, and domain thickness—on bundle extension (Figs. 2 and 3). It was found that our results are consistent with those reported in previous computational studies ^50-52^. When cross-linking density was varied within a range spanning three orders of magnitude, we observed the biphasic dependence of bundle extension on cross-linking density (Fig. 4A). This biphasic dependence emerges due to positive and negative effects of cross-linkers on bundle extension. If cross-linking density is zero or very low, MTs are displaced relatively fast by motors and then push the extending end individually (Fig. 2A). Initially, due to these MTs, a bundle can extend fast, but the extended length is not sustained stably because MTs could be easily buckled by a compressive load (Figs. 2B-D). If there is stronger support from the side of MTs, the buckling can be suppressed (Fig. 3D), but if entire long MTs can be freely bent, a small load can buckle the MTs that directly push the extending end. If there are cross-linkers, the MTs become less susceptible to compression-induced buckling, because the unit of buckling is changed from entire MTs to part of MTs located between adjacent cross-linking points, according to Eq. 10. With cross-linkers, the curvature of entire MTs slightly increased across the bundle due to interactions between cross-linkers and motors (Fig. 4B), but the curvature of MTs directly pushing the extending end becomes smaller (Fig. 4E), which directly contributes to enhanced bundle extension. However, at the same time, the movement of MTs becomes harder to occur due to many transient connections to other MTs. Motors feel larger resistance from cross-linkers when they attempt to displace MTs by walking toward the minus end of MTs, leading to larger force development on motors and slower walking motions (Figs. 4F, G). In this case, it takes much more time for motors to develop forces on MTs, and bundle extension could be much smaller. Due to these two opposite effects of cross-linkers, bundle extension appears to be maximal at the intermediate cross-linking level. In addition, the stiffness of MTs was found to be another key factor regulating bundle extension. If MTs are stiffer, they are less likely to undergo buckling against the same compressive loads (Figs. 5B-E), so the biphasic dependence of bundle extension on cross-linking level disappears (Fig. 5F), and bundle extension is enhanced (Fig. 5A). However, if there are too many cross-linkers, an increase in the MT stiffness cannot enhance bundle extension significantly because the motions of MTs are too hard to take place.

Our observations are in good agreements with numerical analyses ^61^ and *in-vitro* experiments ^37,62^ which hypothesized that MTs reinforced with cross-linkers are capable of enduring greater compressive loads before experiencing buckling events. Another notable finding was that MT extends the fastest at an optimal range of *R*_X_, following previous observations that the overexpression of tau stimulates neurite outgrowth or can even rescue impaired outgrowth ^63,64^. Admittedly, our model has several limitations. Our model consists of minimal components: MTs with the same polarity, cross-linkers, and dynein-like motors. Previous studies reported mixed MT polarity in axons ^36,50^ or other cytoskeletal structures such as actomyosin cortex surrounding the MT array ^35^. It is expected that the inclusion of these antagonistic factors in the model would result in slower bundle extension. Moreover, in our study, only a limited set of parameters were varied, and their combinatorial effects were not fully explored although we showed how interplay between cross-linkers and MT stiffness mediated bundle extension. Noting that mechanical factors for neuron elongation are pushing forces from MT bundles along the axon and pulling forces from actin filaments at the growth cone ^28^, we next aim to incorporate actin dynamics as an additional driver of neurite elongation.

## CONCLUSION

In this study, we demonstrated how the extension of MT bundles, which resemble elongating axon structures, is regulated by cross-linkers and the stiffness of MTs. Cross-linkers enabled relatively soft MTs to extend the bundle against a mechanical resistance by preventing MTs from being buckled by compressive loads. At the same time, cross-linkers reduced bundle extension by restricting MT movements. This led to the emergence of maximal bundle extension at intermediate cross-linking level. However, when MTs were sufficiently stiff, the bundle extension was maximal at a wide range of low cross-linking density because the cross-linker-induced enforcement of MTs did not play a significant role in bundle extension. Results on MT mechanics during extension provide insights into understanding the intrinsic mechanism of axonal outgrowth. In future studies, we will incorporate the dynamic behaviors of MTs to probe interplay between MT mechanics and dynamics for bundle extension and will incorporate actomyosin cortex around MTs to systematically probe competitions between contractile forces generated by the cortex and extensile forces exerted by MTs.

## SUPPLEMENTARY FIGURES

**Fig. S1.**
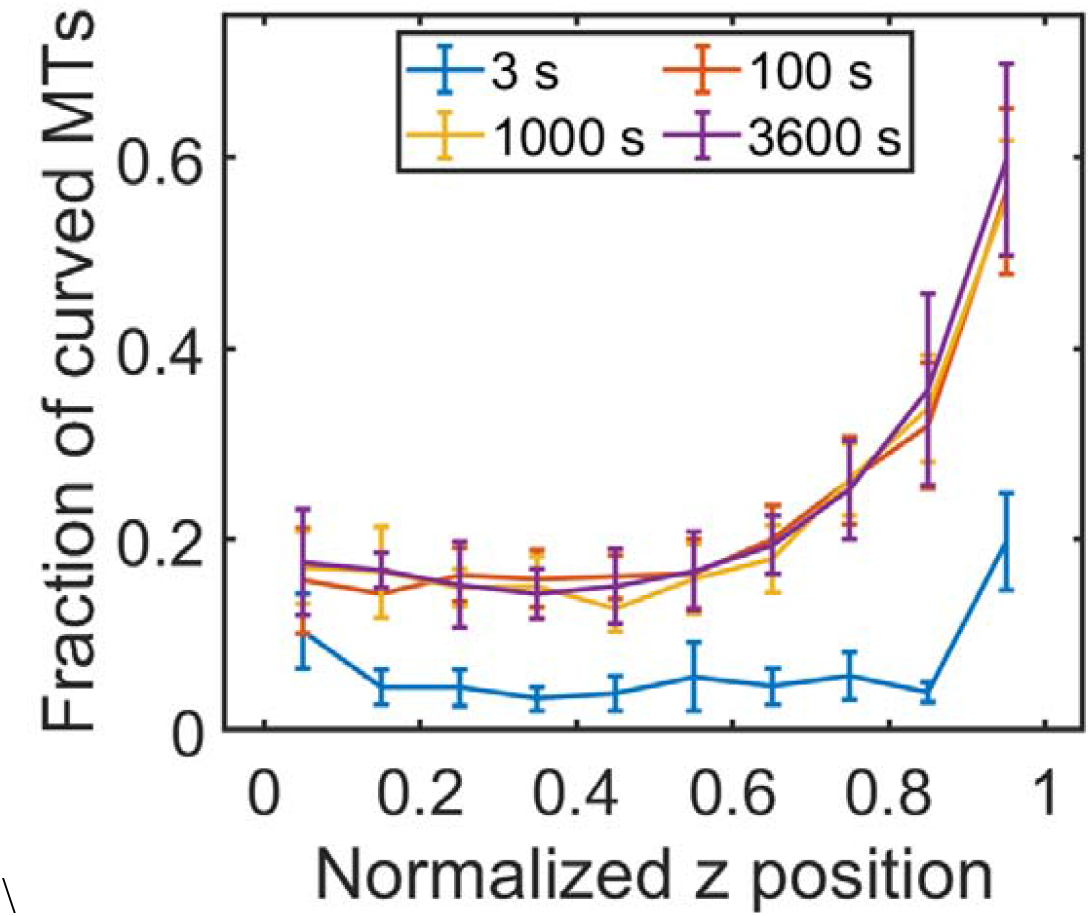
Spatial analysis of the fraction of curved microtubules (MTs) without cross-linkers. 10 simulations used for Fig. 2 were analyzed to estimate how many MTs were buckled in each axial position. The z positions of MT elements were normalized by instantaneous domain length (*L*_dom_). The fraction of MT segments exhibiting curvature greater than 0.2 rad/µm was calculated at 4 time points.

**Fig. S2.**
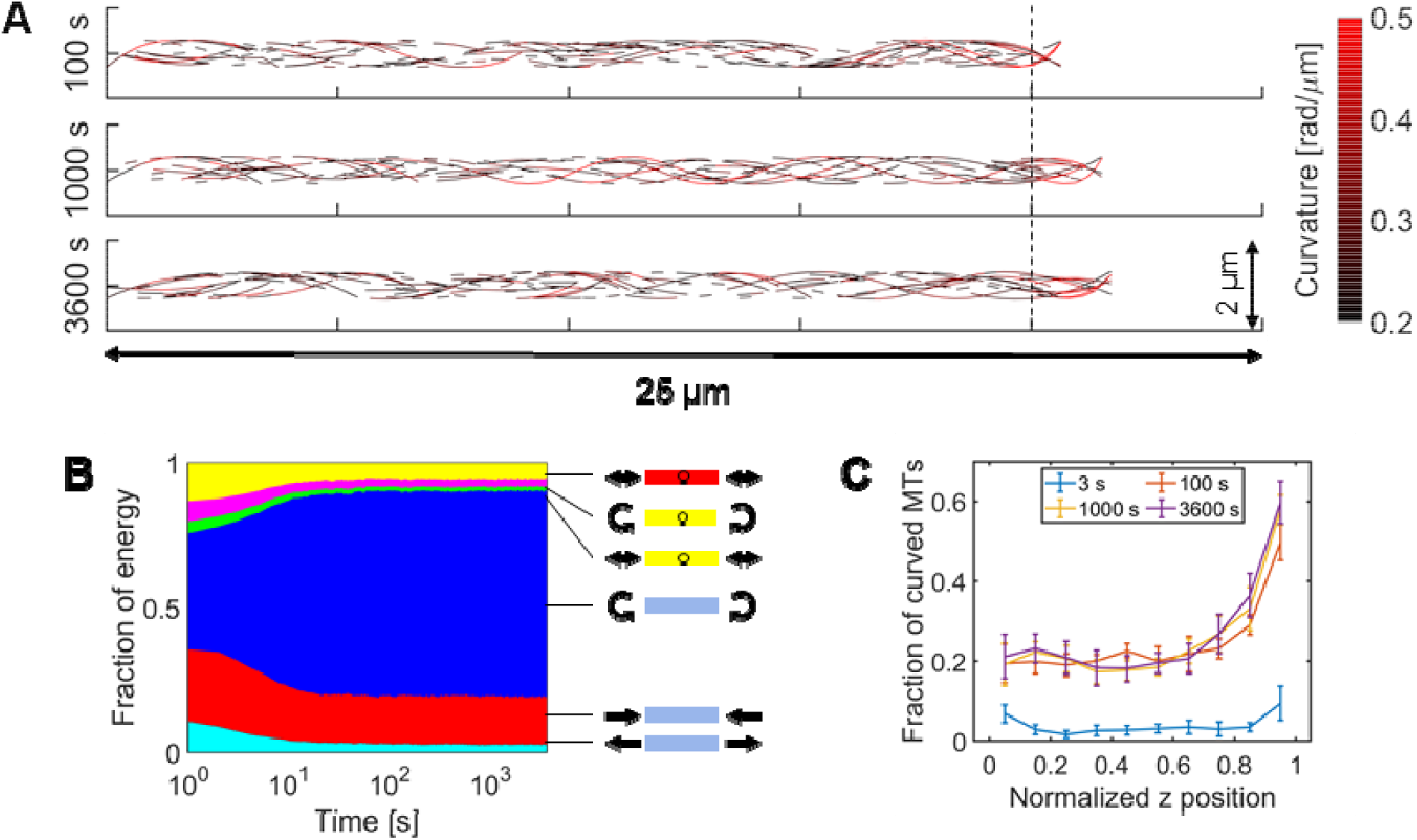
Bundle extension with cross-linkers. (A) Snapshots at three time points showing a fraction of microtubules (MTs) with curvature higher than 0.2 rad/µm. With *R*_X_ = 0.25 µm^-1^, only MTs were visualized in these snapshots, for clarity. 10 simulations were performed for 1 hr under this condition. Even with cross-linkers, MTs located near the extending end on the right are more curved than others. (B) The fraction of six types of energies over time: tensile (cyan), compressive (red), and bending (blue) energies of MTs, the extensional (green) and bending (magenta) energies of cross-linkers, and the extensional energy of motors (yellow). (C) The fraction of curved MTs depending on axial (*z*) positions at 4 time points.

**Fig. S3.**
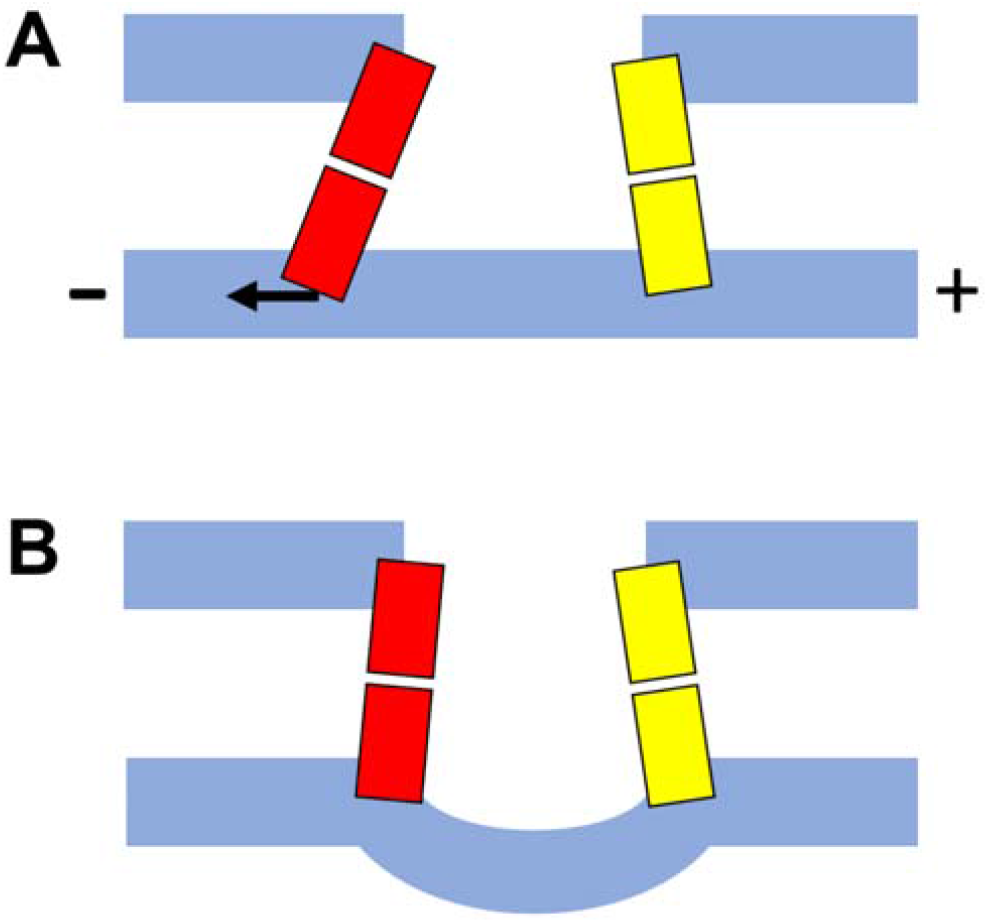
Buckling of microtubules (MTs) occurring when MT translocation is restricted by cross-linkers. (A) A motor (red) walks toward the minus end. As a result, the bottom MT is translocated to the right. If this MT is connected to other MT via cross-linkers (yellow), translocation can be frustrated beyond cross-linking points. Then, the MT can be curved as shown in (B). Note that buckling of MTs requires a compressive force larger than a maximum force that a single motor can generate, so multiple motors are involved with the buckling of a single MT.

**Fig. S4.**
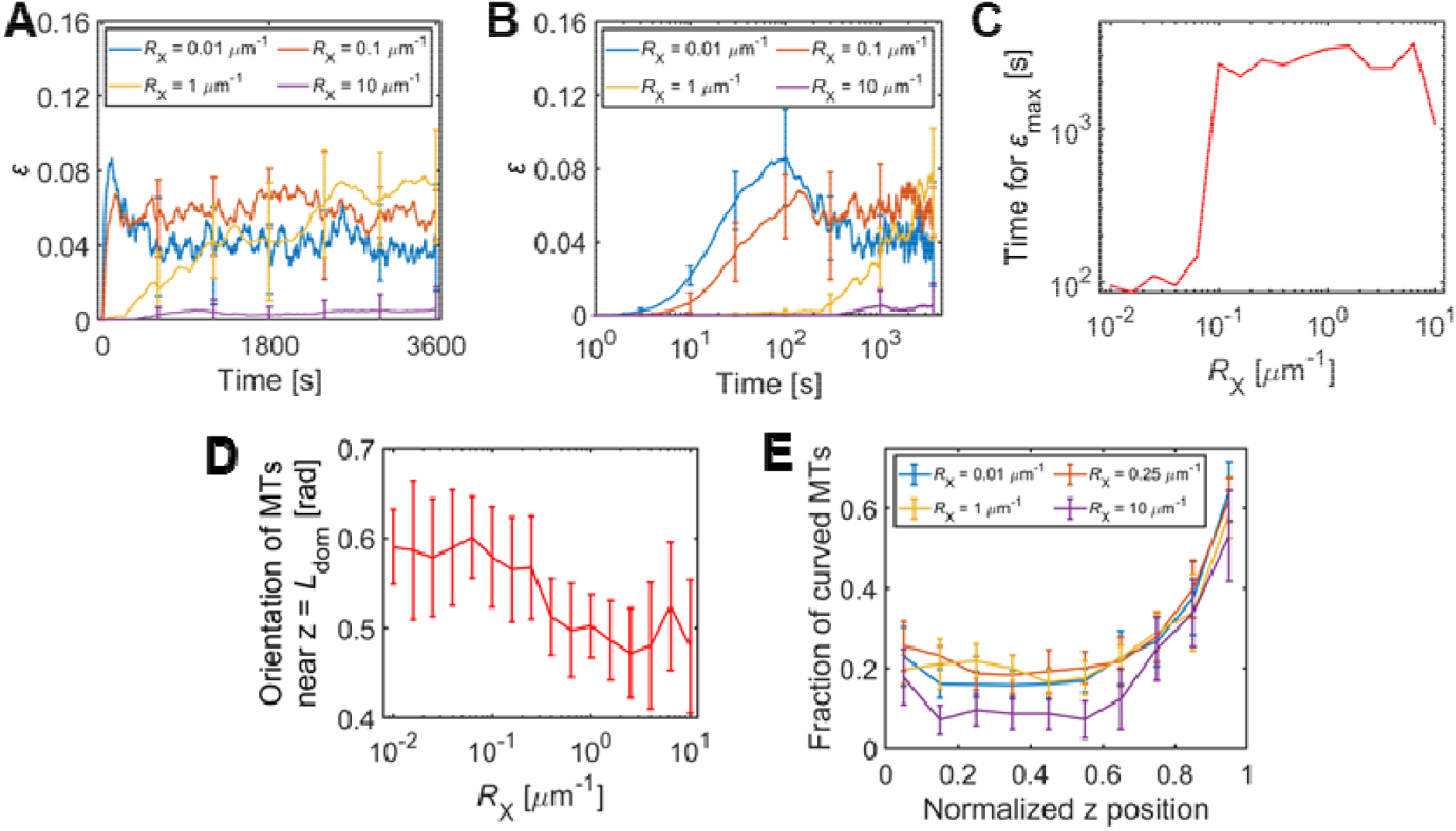
Bundle extension with various cross-linking density (*R*_X_). Simulations shown in Fig. 4 were used for additional analyses. (A, B) Time evolution of normal strain (*ε*) indicating the level of bundle extension with 4 different *R*_X_. Time is shown in the linear scale (A) or the log scale (B). (C) Time for reaching maximal normal strain (*ε*_max_) with various *R*_X_. (D) The orientation of MTs located near the extending end. (E) The fraction of curved MTs depending on axial (*z*) positions depending on *R*_X_.

**Fig. S5.**
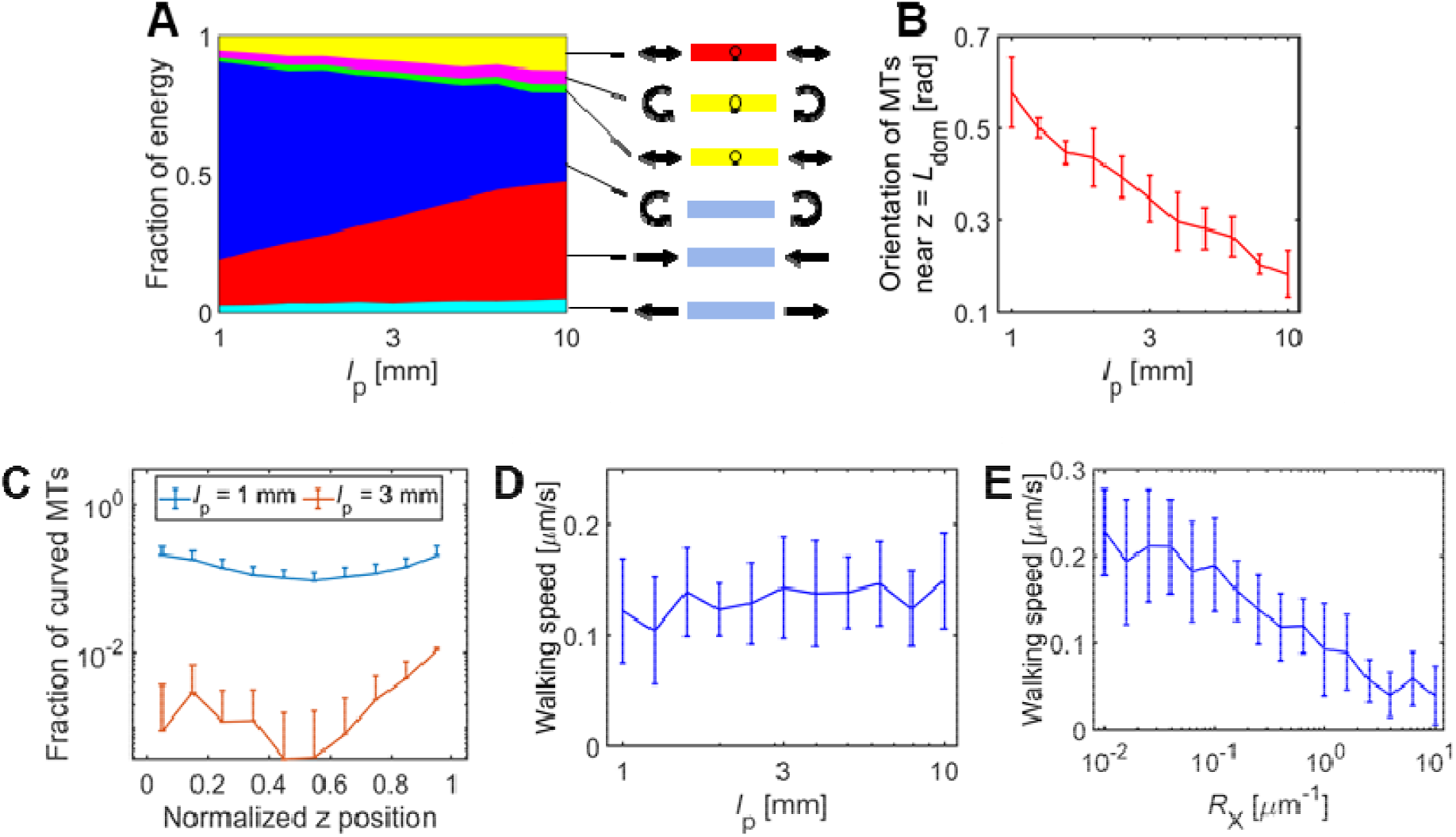
Bundle extension depending on the persistence length of microtubules (*l*_p_). (A) The fraction of six types of energies with different *l*_p_: tensile (cyan), compressive (red), and bending (blue) energies of MTs, the extensional (green) and bending (magenta) energies of cross-linkers, and the extensional energy of motors (yellow). (B) The fraction of curved MTs depending on axial (*z*) positions depending on *l*_p_. (C) The fraction of curved MTs depending on axial (*z*) positions depending on *l*_p_. (D) Average walking speed of motors as a function of *l*_p_. (E) Average walking speed of motors with *l*_p_ = 3 mm and various *R*_X_.

## MOVIE CAPTIONS

**Movie S1. Bundle extension when cross-linking density is very low (*R***_**X**_ **= 0.01 µm**^**-1**^). The curvature of MTs is visualized via color scaling. Motors and cross-linkers are not visualized for clarity. The vertical and horizontal dimensions of the visualized space are 2 μm and 25 μm, respectively.

**Movie S2. Bundle extension with intermediate cross-linking density (*R***_**X**_ **= 0.25 µm**^**-1**^). Only microtubules are visualized with their curvature represented via color scaling. The vertical and horizontal dimensions of the visualized space are 2 μm and 25 μm, respectively.

**Movie S3. Bundle extension with high cross-linking density (*R***_**X**_ **= 10 µm**^**-1**^). The curvature of microtubules is indicated by color scaling without motors and cross-linkers. The vertical and horizontal dimensions of the visualized space are 2 μm and 25 μm, respectively.

**Movie S4. Bundle extension with intermediate persistence length of MTs (*l***_**p**_ **= 3 mm)**. Only microtubules are visualized with their curvature indicated via color scaling, without motors and cross-linkers. The vertical and horizontal dimensions of the visualized space are 2 μm and 25 μm, respectively.

**Movie S5. Bundle extension with high persistence length (*l***_**p**_ **= 10 mm)**. The curvature of microtubules is shown via color scaling. Motors and cross-linkers are not visualized for clarity. The vertical and horizontal dimensions of the visualized space are 2 μm and 25 μm, respectively.

## REFERENCES

1 Polleux, F. & Snider, W. Initiating and growing an axon. Cold Spring Harb Perspect Biol 2, a001925 (2010).

2 Takei, Y., Teng, J., Harada, A. & Hirokawa, N. Defects in axonal elongation and neuronal migration in mice with disrupted tau and map1b genes. J Cell Biol 150, 989–1000 (2000).

3 Suter, D. M. & Miller, K. E. The emerging role of forces in axonal elongation. Prog Neurobiol 94, 91–101 (2011).

4 Lu, W. & Gelfand, V. I. Moonlighting motors: kinesin, dynein, and cell polarity. Trends Cell Biol 27, 505–514 (2017).

5 Chia, J. X., Efimova, N. & Svitkina, T. M. Neurite outgrowth is driven by actin polymerization even in the presence of actin polymerization inhibitors. Mol Biol Cell 27, 3695–3704 (2016).

6 Borisy, G. et al. Microtubules: 50 years on from the discovery of tubulin. Nat Rev Mol Cell Biol 17, 322–328 (2016).

7 Yu, W. & Baas, P. W. Changes in microtubule number and length during axon differentiation. J Neurosci 14, 2818–2829 (1994).

8 Conde, C. & Cáceres, A. Microtubule assembly, organization and dynamics in axons and dendrites. Nat Rev Neurosci 10, 319–332 (2009).

9 Eckel, B. D., Cruz, R., Craig, E. M. & Baas, P. W. Microtubule polarity flaws as a treatable driver of neurodegeneration. Brain Res Bull 192, 208–215 (2023).

10 Rao, A. N. et al. Cytoplasmic dynein transports axonal microtubules in a polarity-sorting manner. Cell Rep 19, 2210–2219 (2017).

11 Athamneh, A. I. M. et al. Neurite elongation is highly correlated with bulk forward translocation of microtubules. Sci Rep 7, 7292 (2017).

12 Hirokawa, N., Noda, Y., Tanaka, Y. & Niwa, S. Kinesin superfamily motor proteins and intracellular transport. Nat Rev Mol Cell Biol 10, 682–696 (2009).

13 Bhabha, G., Johnson, G. T., Schroeder, C. M. & Vale, R. D. How dynein moves along microtubules. Trends Biochem Sci 41, 94–105 (2016).

14 Dehmelt, L. & Halpain, S. The MAP2/Tau family of microtubule-associated proteins. Genome Biol 6, 204 (2005).

15 Lewis, S. A., Ivanov, I. E., Lee, G. H. & Cowan, N. J. Organization of microtubules in dendrites and axons is determined by a short hydrophobic zipper in microtubule-associated proteins MAP2 and tau. Nature 342, 498–505 (1989).

16 Kosik, K. S. & Finch, E. A. MAP2 and tau segregate into dendritic and axonal domains after the elaboration of morphologically distinct neurites: an immunocytochemical study of cultured rat cerebrum. J Neurosci 7, 3142–3153 (1987).

17 Ahmad, F. J. et al. Motor proteins regulate force interactions between microtubules and microfilaments in the axon. Nat Cell Biol 2, 276–280 (2000).

18 Dehmelt, L., Nalbant, P., Steffen, W. & Halpain, S. A microtubule-based, dynein-dependent force induces local cell protrusions: Implications for neurite initiation. Brain Cell Biol 35, 39–56 (2006).

19 Roossien, D. H., Lamoureux, P. & Miller, K. E. Cytoplasmic dynein pushes the cytoskeletal meshwork forward during axonal elongation. J Cell Sci 127, 3593–3602 (2014).

20 McElmurry, K. et al. Dynein-mediated microtubule translocation powering neurite outgrowth in chick and Aplysia neurons requires microtubule assembly. J Cell Sci 133, jcs232983 (2020).

21 Ahmad, F. J., Joshi, H. C., Centonze, V. E. & Baas, P. W. Inhibition of microtubule nucleation at the neuronal centrosome compromises axon growth. Neuron 12, 271–280 (1994).

22 Rochlin, M. W., Wickline, K. M. & Bridgman, P. C. Microtubule stability decreases axon elongation but not axoplasm production. J Neurosci 16, 3236–3246 (1996).

23 Tang-Schomer, M. D., Patel, A. R., Baas, P. W. & Smith, D. H. Mechanical breaking of microtubules in axons during dynamic stretch injury underlies delayed elasticity, microtubule disassembly, and axon degeneration. FASEB J 24, 1401–1410 (2010).

24 LaMonte, B. H. et al. Disruption of dynein/dynactin inhibits axonal transport in motor neurons causing late-onset progressive degeneration. Neuron 34, 715–727 (2002).

25 Sainath, R. & Gallo, G. The dynein inhibitor Ciliobrevin D inhibits the bidirectional transport of organelles along sensory axons and impairs NGF-mediated regulation of growth cones and axon branches. Dev Neurobiol 75, 757–777 (2015).

26 Lamoureux, P., Buxbaum, R. E. & Heidemann, S. R. Direct evidence that growth cones pull. Nature 340, 159–162 (1989).

27 Miller, K. E. & Suter, D. M. An integrated cytoskeletal model of neurite outgrowth. Front Cell Neurosci 12, 447 (2018).

28 Letourneau, P. C., Shattuck, T. A. & Ressler, A. H. “Pull” and “push” in neurite elongation: observations on the effects of different concentrations of cytochalasin B and taxol. Cell Motil Cytoskeleton 8, 193–209 (1987).

29 Dubey, S. et al. The axonal actin-spectrin lattice acts as a tension buffering shock absorber. Elife 9, e51772 (2020).

30 Letourneau, P. C. Actin in axons: stable scaffolds and dynamic filaments. Results Probl Cell Differ 48, 65–90 (2009).

31 Malacrida, A., Meregalli, C., Rodriguez-Menendez, V. & Nicolini, G. Chemotherapy-induced peripheral neuropathy and changes in cytoskeleton. Int J Mol Sci 20, 2287 (2019).

32 Xu, K., Zhong, G. & Zhuang, X. Actin, spectrin, and associated proteins form a periodic cytoskeletal structure in axons. Science 339, 452–456 (2013).

33 Kubo, T. et al. Myosin IIA is required for neurite outgrowth inhibition produced by repulsive guidance molecule. J Neurochem 105, 113–126 (2008).

34 Wang, Y. et al. Myosin IIA-related actomyosin contractility mediates oxidative stress-induced neuronal apoptosis. Front Mol Neurosci 10, 75 (2017).

35 de Rooij, R., Miller, K. E. & Kuhl, E. Modeling molecular mechanisms in the axon. Comput Mech 59, 523–537 (2017).

36 Jakobs, M. A. H., Zemel, A. & Franze, K. Unrestrained growth of correctly oriented microtubules instructs axonal microtubule orientation. Elife 11, e77608 (2022).

37 Brangwynne, C. P. et al. Microtubules can bear enhanced compressive loads in living cells because of lateral reinforcement. J Cell Biol 173, 733–741 (2006).

38 Odde, D. J., Ma, L., Briggs, A. H., DeMarco, A. & Kirschner, M. W. Microtubule bending and breaking in living fibroblast cells. J Cell Sci 112, 3283–3288 (1999).

39 Heidemann, S. R., Kaech, S., Buxbaum, R. E. & Matus, A. Direct observations of the mechanical behaviors of the cytoskeleton in living fibroblasts. J Cell Biol 145, 109–122 (1999).

40 Wang, N. et al. Mechanical behavior in living cells consistent with the tensegrity model. Proc Natl Acad Sci U S A 98, 7765–7770 (2001).

41 Tanaka, E. M. & Kirschner, M. W. Microtubule behavior in the growth cones of living neurons during axon elongation. J Cell Biol 115, 345–363 (1991).

42 Schaefer, A. W., Kabir, N. & Forscher, P. Filopodia and actin arcs guide the assembly and transport of two populations of microtubules with unique dynamic parameters in neuronal growth cones. J Cell Biol 158, 139–152 (2002).

43 Underhill, P. T. & Doyle, P. S. On the coarse-graining of polymers into bead-spring chains. J Non-Newton Fluid Mech 122, 3–31 (2004).

44 Clift, R., Grace, J. R. & Weber, M. E. Bubbles, drops, and particles. (2005).

45 Isambert, H. et al. Flexibility of actin filaments derived from thermal fluctuations. Effect of bound nucleotide, phalloidin, and muscle regulatory proteins. J Biol Chem 270, 11437–11444 (1995).

46 Jung, W., P Murrell, M. & Kim, T. F-actin cross-linking enhances the stability of force generation in disordered actomyosin networks. Comput Part Mech 2, 317–327 (2015).

47 Bell, G. I. Models for the specific adhesion of cells to cells. Science 200, 618–627 (1978).

48 Monzon, G. A., Scharrel, L., Santen, L. & Diez, S. Activation of mammalian cytoplasmic dynein in multimotor motility assays. J Cell Sci 132, jcs220079 (2018).

49 Memet, E. et al. Microtubules soften due to cross-sectional flattening. Elife 7, e34695 (2018).

50 Jakobs, M. A. H., Franze, K. & Zemel, A. Mechanical regulation of neurite polarization and growth: a computational study. Biophys J 118, 1914–1920 (2020).

51 Jakobs, M., Franze, K. & Zemel, A. Force generation by molecular-motor-powered microtubule bundles; implications for neuronal polarization and growth. Front Cell Neurosci 9, 441 (2015).

52 de Rooij, R., Kuhl, E. & Miller, K. E. Modeling the axon as an active partner with the growth cone in axonal elongation. Biophys J 115, 1783–1795 (2018).

53 Voelzmann, A., Hahn, I., Pearce, S. P., Sánchez-Soriano, N. & Prokop, A. A conceptual view at microtubule plus end dynamics in neuronal axons. Brain Res Bull 126, 226–237 (2016).

54 Weingarten, M. D., Lockwood, A. H., Hwo, S. Y. & Kirschner, M. W. A protein factor essential for microtubule assembly. Proc Natl Acad Sci U S A 72, 1858–1862 (1975).

55 Chen, J., Kanai, Y., Cowan, N. J. & Hirokawa, N. Projection domains of MAP2 and tau determine spacings between microtubules in dendrites and axons. Nature 360, 674–677 (1992).

56 Lansky, Z. et al. Diffusible crosslinkers generate directed forces in microtubule networks. Cell 160, 1159–1168 (2015).

57 Ahmad, F. J., Echeverri, C. J., Vallee, R. B. & Baas, P. W. Cytoplasmic dynein and dynactin are required for the transport of microtubules into the axon. J Cell Biol 140, 391–401 (1998).

58 Esmaeli-Azad, B., McCarty, J. H. & Feinstein, S. C. Sense and antisense transfection analysis of tau function: tau influences net microtubule assembly, neurite outgrowth and neuritic stability. J Cell Sci 107, 869–879 (1994).

59 Tofangchi, A., Fan, A. & Saif, M. T. A. Mechanism of axonal contractility in embryonic drosophila motor neurons in vivo. Biophys J 111, 1519–1527 (2016).

60 Soheilypour, M., Peyro, M., Peter, S. J. & Mofrad, M. R. K. Buckling behavior of individual and bundled microtubules. Biophys J 108, 1718–1726 (2015).

61 Brodland, G. W. & Gordon, R. Intermediate filaments may prevent buckling of compressively loaded microtubules. J Biomech Eng 112, 319–321 (1990).

62 Jin, M. Z. & Ru, C. Q. Localized buckling of a microtubule surrounded by randomly distributed cross linkers. Phys Rev E Stat Nonlin Soft Matter Phys 88, 012701 (2013).

63 Yoshizaki, C., Tsukane, M. & Yamauchi, T. Overexpression of tau leads to the stimulation of neurite outgrowth, the activation of caspase 3 activity, and accumulation and phosphorylation of tau in neuroblastoma cells on cAMP treatment. Neurosci Res 49, 363–371 (2004).

64 Zuo, Y. C. et al. Overexpression of Tau rescues Nogo-66-induced neurite outgrowth inhibition in vitro. Neurosci Bull 32, 577–584 (2016).

